# RNA-seq analysis of laser micro-dissected Arabidopsis thaliana leaf epidermis, mesophyll and vasculature defines tissue-specific transcriptional responses to multiple stress treatments

**DOI:** 10.1101/2020.11.01.364257

**Authors:** Oliver Berkowitz, Yue Xu, Yan Wang, Lim Chee Liew, Yanqiao Zhu, Mathew G. Lewsey, James Whelan

**Affiliations:** Department of Animal, Plant and Soil Science, ARC Centre of Excellence in Plant Energy Biology, La Trobe University, Bundoora, Victoria 3086, Australia; Department of Animal, Plant and Soil Science, La Trobe University, Bundoora, Victoria 3086, Australia

## Abstract

Acclimation of plants to adverse environmental conditions requires the coordination of gene expression and signalling pathways between tissues and cell types. As the energy and carbon capturing organs, leaves are significantly affected by abiotic and biotic stresses. However, tissue- or cell type-specific analyses of stress responses have largely focussed on the Arabidopsis root. Here, we comparatively explore the transcriptomes of three leaf tissues (epidermis, mesophyll, vasculature) after induction of diverse stress pathways by chemical stimuli (antimycin A, 3-amino-1,2,4-triazole, methyl viologen, salicylic acid) and UV light in Arabidopsis. Profiling stimuli-dependent changes after treatments revealed an overall reduction in the tissue-specific expression of genes, with only a limited number gaining or changing their tissue-specificity. We find no evidence of a common stress response, with only a few genes responsive to two or more treatments in the analysed tissues. However, differentially expressed genes overlap across tissues for individual treatments. Further analyses provided evidence for an interaction of auxin and ethylene that mediates retrograde signalling during mitochondrial dysfunction specifically in the epidermis, and a gene regulatory network defined the hierarchy of interactions. Taken together, we generated an extensive reference data set and results enable the tailoring of the tissue-specific engineering of stress tolerant plants.

## INTRODUCTION

Crop losses due to abiotic and biotic stresses substantially reduce agricultural production each year, which impacts global food security in a changing climate. Consequently, improving tolerance to adverse conditions is an important goal for plant scientists and breeders who must sustain increases in crop yields over the coming decades (Bailey-Serres et al., 2019; Foley et al., 2011). Conventional breeding methods are laborious and time consuming, limiting the speedy development of improved genetic material (Hickey et al., 2019). However, stacking of traits through molecular genetics using known, underlying target genes is able to recapitulate such decades-long breeding achievements or even domestication within a few years (Rodríguez-Leal et al., 2017; Zsögön et al., 2018). Although this shows the potential of advances in gene editing, application of such technologies is hampered by identification of suitable target genes or combinations that lead to stress tolerance. One limitation is incomplete understanding of the complex interplay of tolerance mechanisms and their signalling pathways that occur in the field (Lamers et al., 2020; Zandalinas et al., 2019). Furthermore, stresses often impact plant tissues to varying degrees and concomitantly stress responses have often tissue-specific components (Long, 2011). However, most analyses of plant stress responses have been performed at a whole-organ level. This may give ambiguous results as cells can be at different stages on the response trajectory or have distinct responses due to their function. Therefore observed gene expression levels may result in inappropriate conclusions based on Simpson’s paradox (Simpson, 1951; Trapnell, 2015). Hence, molecular approaches on a tissue- or cell-specific level show great potential to identify novel or more appropriate targets and pathways for the improvement of stress tolerance. The identification of such gate-keeper cell types provides targets to tailor plant engineering for tolerance (Henderson and Gilliham, 2015).

Advances in methodology have made it possible to analyse transcriptomes on the tissue, cell-type and single-cell level. The most widely used techniques are laser capture microdissection (LCM), fluorescence-activated cell sorting (FACS), isolation of nuclei tagged in specific cell types (INTACT) and droplet based single-cell methods (for review see Bailey-Serres, 2013; McFaline-Figueroa et al., 2020). Major breakthroughs for LCM were the global expression analysis of epidermal and vasculature tissues in maize (Nakazono et al., 2003) and the generation of a transcriptome atlas across 40 cell types of the shoot and root in rice (Jiao et al., 2009). INTACT based methodology (Zanetti et al., 2005) profiled gene expression during embryo development (Palovaara et al., 2017), the response of 21 cell types to hypoxia (Mustroph et al., 2009) as well as chromatin modification in root epidermal cells (Deal and Henikoff, 2010). FACS and single-cell based approaches have mostly focussed on the development (Denyer et al., 2019; Jean-Baptiste et al., 2019; Ryu et al., 2019; Shulse et al., 2019; Zhang et al., 2019) and stress responses (Dinneny et al., 2008; Geng et al., 2013; Gifford et al., 2008; Iyer-Pascuzzi et al., 2011; Jean-Baptiste et al., 2019; Long et al., 2010) of the Arabidopsis root. FACS has also successfully been used to survey gene expression in guard cells of Arabidopsis leaves (Gardner et al., 2008; Lee et al., 2019).

These methods all have their specific advantages and disadvantages with respect to their applicability to plant species, organs, preparation of samples, their impact on gene expression, depth, resolution and cost (Bailey-Serres, 2013; McFaline-Figueroa et al., 2020). LCM is independent of genotype or species and relies only on the histological differentiation of cell types but requires laborious tissue fixation and sectioning, limiting the number of experiments and cell types. FACS and INTACT can isolate many cell-types and are reasonably high-throughput. However, both rely on transgenic plants expressing tagged (fluorescent) proteins in the targeted cell types and autofluorescent cells types, e.g. from the leaf, are difficult to isolate (Galbraith, 2007). A limitation of FACS, INTACT and single-cell methods is the necessary protoplasting of tissues which biases the capturing of cell types from a tissue and also introduces changes in gene expression to a yet unknown extent. Single-cell methods provide the highest resolution, allowing characterisation of dynamic changes in cell types and states. Their analysis relies however on specific marker genes for the identification cell types. Changes in the expression profiles of these genes under varying conditions however impacts cell-type assignment, and detection of cell type—specific differences becomes difficult when stress responses overwhelm cell-specific signatures (Jean-Baptiste et al., 2019).

Studies on transcriptional responses of tissue- or cell type-specific to stress have focussed on Arabidopsis roots (Dinneny et al., 2008; Geng et al., 2013; Gifford et al., 2008; Iyer-Pascuzzi et al., 2011; Jean-Baptiste et al., 2019; Rich-Griffin et al., 2020). Similar studies on leaves are lacking, although leaves are the primary energy provider through photosynthesis and a main target of many abiotic and biotic stresses. The three major tissues of Arabidopsis leaves are the epidermis, mesophyll and vasculature and each contributes to varying degrees to the overall mature leaf (epidermis: 36 %, mesophyll: 36%, vasculature: 28%) (Pyke and López-Juez, 1999). These tissues perform a number of classical functions, but more recent research also points to other roles. The epidermis forms a protective layer and regulates gas and water exchange via stomata (Javelle et al., 2011), but also drives or restricts the growth of the whole organ via brassinosteroid perception (Savaldi-Goldstein et al., 2007) or coordinates auxin-dependent shade avoidance (Procko et al., 2016). While the mesophyll is the main site for photosynthesis and carbon fixation, chloroplasts are also the start and hub of signalling pathways important for the whole plant (Chan et al., 2016). The vasculature, consisting of the phloem and xylem, forms a network throughout the leaf for water, nutrient and solute transport. In addition to this, the phloem and xylem also play a role in long-distance signalling between the root and shoot via signalling components which include microRNAs and hormones (Lucas et al., 2013) and integration of rapid systemic signals such as ROS (Zandalinas et al., 2020). These specialisation in cell types and functional diversity have also led to varying organelle organisations. Consequently, the number of chloroplasts and mitochondria contrasts greatly in the different leaf tissues (Barton et al., 2016; Cayla et al., 2015; Logan, 2006) as does their association (Islam et al., 2009). These adaptations of organelles to overall tissue function are also impacting stress responses (Armstrong et al., 2006), especially as recent evidence suggest that these organelles also perform a role as signalling hubs and stress sensors (Crawford et al., 2017; Kleine and Leister, 2016; Wang et al., 2020).

In this study we analysed the transcriptomic response of the leaf epidermis, mesophyll and vasculature to five treatments selected to induce varying stress signalling pathways by laser microdissection and RNA-seq (LCM-seq). This makes available a compendium of the cell-type specific expression of genes, their changes following stress induction and conserved tissue-specific marker genes. While tissue-specificity in gene expression after treatments is in generally reduced, tissues share transcriptional responses to individual treatments. In addition, we provide evidence for the regulation of mitochondrial signalling by auxin and ethylene in the epidermis.

## RESULTS

### Laser capture microdissection followed by RNAseq enables tissue- and treatment specific analysis of transcriptomic responses

To enable the analysis of tissue-specific changes in gene expression after chemical, hormone or other abiotic or biotic stress treatments, we established methodology using laser capture microdissection (LCM) followed by next-generation sequencing (LCM-seq). This methodology was applied to plants treated with four chemicals and ultraviolet C (UV-C) light by isolation of RNA from the three major leaf tissues of Arabidopsis, i.e. epidermis, mesophyll and vasculature (Figure 1A). These treatments were chosen for their ability to trigger a range of specific responses dependent on their mode of action in plants: 1) antimycin A (AA) is an inhibitor of the mitochondrial electron transport chain leading to mitochondrial dysfunction (Labs et al., 2016), 2) 3-amino-1,2,4-triazol (3AT) inhibits several metabolic reactions including catalase and carotenoid biosynthesis (Feierabend and Winkelhüsener, 1982), 3) methyl viologen (MV, also known as paraquat) inhibits photosynthesis by accepting electrons from photosystem I and leads to superoxide (O_2_^•-^) production in chloroplasts (Fuerst and Norman, 1991), 4) salicylic acid is a plant hormone activating biotic stress signalling cascades (Vlot et al., 2009), and 5) ultraviolet light (UV) damages DNA, protein and membrane structure and induces ROS accumulation (Müller-Xing et al., 2014). Therefore, these treatments were expected to have varying degrees of impact on the tissues analysed to result in distinct and varying transcriptional responses. The LCM method yielded on average 1 ng total RNA as the input for a low input library generation protocol for subsequent RNA-seq. This gave an average of 6.7 M reads of high quality (Q30 > 90%) per biological replicate and pseudo-aligning to the Araport 11 annotated transcripts.

**Figure 1.**
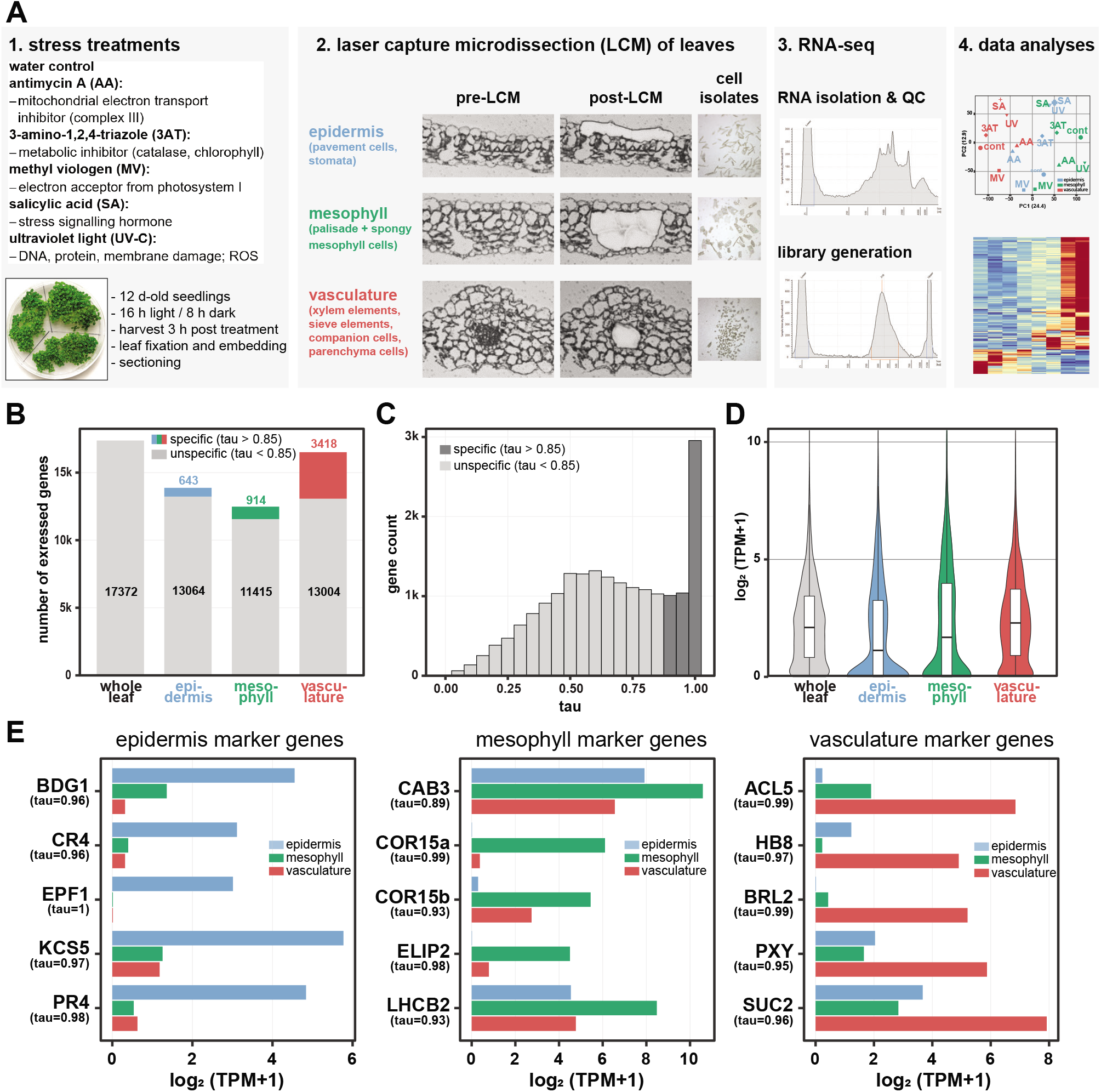
Overview of experimental setup and tissue-specific gene expression under control conditions. **(A)** Illustration of the experimental setup for stress treatments and laser capture microdissection for subsequent RNAseq analyses. Arabidopsis seedlings (accession Col-0, grown in vitro for 12 days) were treated by spraying with antimycin A (AA; 50 μM), 3-amino-1,2,4-triazole (3AT; 10 mM), methyl viologen (MV; 1 mM) and salicylic acid (SA; 2 mM), and by exposure to ultraviolet light (UV-C; 1.5 W/m^2^) for 20 min. For controls plants were sprayed with water. Leaves were harvested 3 h post treatment, fixed, paraffin embedded and sectioned. Epidermis, mesophyll and vasculature were isolated by laser capture microdissection for RNA extraction and library generation for next-generation sequencing. RNA-seq data was then analysed for (differential) gene expression. **(B)** Expression of genes under control conditions in three leaf tissues (epidermis, blue; mesophyll, green; vasculature, red) was quantified on a transcripts per million (TPM) basis and compared to a RNA-seq data set for whole leaf. Genes were called as expressed in either the whole leaf or the different tissues using a TPM ≥ 1 cut-off (Supplemental Table 1). The bar chart represents the number of genes expressed in the three tissues and the whole leaf for comparison. Tissue-specificity was defined using the tau measure (Kryuchkova-Mostacci and Robinson-Rechavi, 2016). The coloured sections of the bars indicate the number of genes specific for each tissues with a tau > 0.85. **(C)** Distribution of tau values for genes detected across the three tissues. Dark grey bars highlight tissue-specific values of tau ≥ 0.85. **(D)** Violin plots showing the distribution of expression for genes with tissue-specific expression (tau ≥ 0.85) and of genes in the whole leaf. **(E)** Expression of five known tissue-specific marker genes for each of the three tissues analysed. All marker genes are highly specific with tau values well above the cut-off of 0.85 used to define tissue-specificity.

### Substantial variation of gene expression in leaf tissues

We first examined how tissue-specific gene expression profiles differ from profiles of bulk organs. To do so we compared the gene expression in the three dissected leaf tissues of plants grown under control conditions with an RNAseq dataset for whole leaves (Meng et al., 2019). The number of expressed genes detectable, using a transcripts per million (TPM) cut-off of TPM ≥ 1, in epidermis (13,707 genes), mesophyll (12,329 genes) and vasculature (16,422 genes) was lower than in whole leaves (17,297 genes) (Figure 1B; Supplemental Table ST1). Genes preferentially expressed in one tissue were identified using the tau metric with a cut-off of tau > 0.85, which provides a robust measure of tissue-specificity (Kryuchkova-Mostacci and Robinson-Rechavi, 2016). The highest number of tissue-specific genes was found in the vasculature (3,418 genes), followed by the mesophyll (914 genes) and the epidermis (643 genes) (Figure 1B; Supplemental Table ST1). These numbers likely reflect the complexity of the isolated cell types (Figure 1A), with the epidermis and mesophyll having less diverse cell types (Pyke and López-Juez, 1999). These proportions are concordant with an earlier report that used the alternative INTACT approach to isolate specific tissues (Mustroph et al., 2009).The vasculature not only transports nutrient and water, but is also an information ‘highway’ and sensory tissue with high transcriptional responsiveness (Long, 2011; Lough and Lucas, 2006). Except for the tissue-specific genes (tau > 0.85), most genes had a tau value of around 0.5, indicating their significant expression in more than one tissue (Figure 1C). Compared to the expression of genes in the whole tissue sample, the vasculature-specific genes showed a similar distribution of expression, while epidermis- and mesophyll had a higher number of genes with low expression (Figure 1D).

We next analysed the functions of genes expressed specifically in single tissues to better understand how the genes relate to the known properties of those tissues. A GO term enrichment analysis for the three lists of tissue-specific genes found enriched GO terms relating to lipid and wax metabolism, stomatal development and auxin response for the epidermis-specific genes (Supplemental Figure 1, Supplemental Table 2). This included members of the *3-KETOACYL-COA SYNTHASE* (*KCS*), *GLYCEROL-3-PHOSPHATE SN-2-ACYLTRANSFERASE* (*GPAT*), *ECERIFERUM* (*CER*) and *LONG-CHAIN ACYL-COA SYNTHETASE* (*LACS*) gene families and the MYB DOMAIN PROTEIN (MYB) transcription factors MYB16 and MYB30, all involved in the wax synthesis for epidermal cuticule formation (Kunst and Samuels, 2003; Pollard et al., 2008; Raffaele et al., 2008). The auxin response is also likely associated to epidermal growth involving ethylene (Swarup et al., 2005; Vaseva et al., 2018). For the list of genes showing vasculature-specific expression, enriched GO terms are associated with xylem/phloem development, cell wall biogenesis and mitosis as well as protein synthesis and turn-over (Supplemental Figure 1, Supplemental Table 2). Corresponding genes encode a number of NAC DOMAIN CONTAINING PROTEIN (ANAC) and MYB transcription factors which are key regulators of secondary cell wall biosynthesis, especially in the xylem (Fukuda and Ohashi-Ito, 2019). In addition, *IRREGULAR XYLEM* (*IRX*) genes, *CLAVATA3/ESR-RELATED* (*CLE*) genes, cellulose synthases-encoding genes as well as *PHLOEM INTERCALATED WITH XYLEM* (*PXE*) and *REVOLUTA* (*REV*) were also included in these GO terms. Mitosis associated GO terms with a number of genes encoding cyclins, cyclin-dependent kinases, cellulose synthases and others are also indicating the high rate of cell divisions in the developing vascular tissue (Fukuda and Ohashi-Ito, 2019). The enriched GO terms for the mesophyll-specific gene set were related to a response to hormones, oxidative stress, defence and photosynthesis (Supplemental Figure 1, Supplemental Table 2). Corresponding genes included a number of *ETHYLENE RESPONES FACTOR* (*ERF*) and *WRKY DNA-BINDING PROTEIN* (*WRKY*) genes. Given that the main functions of the mesophyll are related to photosynthesis, a surprisingly low number of enriched GO terms and associated genes for this process were found as specific for this tissue. A possible explanation is that functional chloroplasts also occur in the epidermis (Barton et al., 2016) and vasculature (Cayla et al., 2015), and hence photosynthesis-related genes are not exclusive to the mesophyll.

To further benchmark the identification of tissue-specific genes by our approach, we extracted the expression of five marker genes for each of the three tissues from our LCM-seq data. Tissue specificity of the epidermis and vasculature marker genes was very high, with tau values ranging from 0.95 to 1, corresponding to at least 10-fold higher expression in the respective tissue than the next lowest (Figure 1E). For the mesophyll marker genes COR15a and ELIP2 the tau values were in a similar range. The tau value for the *CHLOROPHYLL A/B BINDING PROTEIN 3* (*CAB3*) was distinctly the lowest among these maker genes (tau = 0.89), a consequence of a significant expression of this gene also in the epidermis and vasculature (Figure 1E). Although genes such as *CAB3* are frequently used as mesophyll marker genes, our data set has identified many genes with higher specificity (Supplemental Table 1). In our list of 914 mesophyll-specific genes (tau > 0.85), 83 % had a greater tau value than *CAB3*, and 60% had a tau > 0.95, i.e. are superior markers.

Taken together, these results confirm that our LCM-seq experiment provides robust tissue-specific gene expression data for three leaf tissues. This also provides some ground truth for single-cell analysis of leaves, for which reference expression data sets and specific marker genes are crucial for cell type identification (McFaline-Figueroa et al., 2020).

### Stress treatments reduce tissue-specific signatures of gene expression

Abiotic and biotic stresses induce substantial changes in gene expression. However, the contribution of distinct leaf tissues to these responses is largely unknown. Using the LCM-seq approach we first analysed if the four different chemical stimuli (3AT, AA, MV, SA) and UV impacted on the tissue-specificity of genes. A principle component analyses of the RNA-seq data for the treatment experiments showed separation by tissue type (Figure 2A; PC1) and treatments (Figure 2A; PC2), indicating differences in the response of the three tissues to the treatments. The response to the five treatments was affected to varying degrees in the three tissues as indicated by changes in the distribution of gene expression levels (Figure 2B). As already observed for the control treatment (Figure 1D), the epidermis and mesophyll had a higher number of genes with low expression level than the vasculature after all treatments, with the latter having a high number of genes with an expression level of above 3 TPM (Figure 2B). Across the treatments there were some specific changes in these distributions. In the epidermis, the distribution trended towards higher TPM values for the AA and 3AT treatments, while for the mesophyll there was a slight shift of low expressed genes to higher expression for MV. The latter might indicate a more pronounced effect of MV on photosynthetic tissues, which would be consistent with its mode of action. In the vasculature treatment with 3AT led to a marked decrease in genes with low expression (Figure 2B). Overall, the pattern for differentially expressed genes (DEGs) varied for each treatment across tissues (Figure 2C, inner circle).

**Figure 2:**
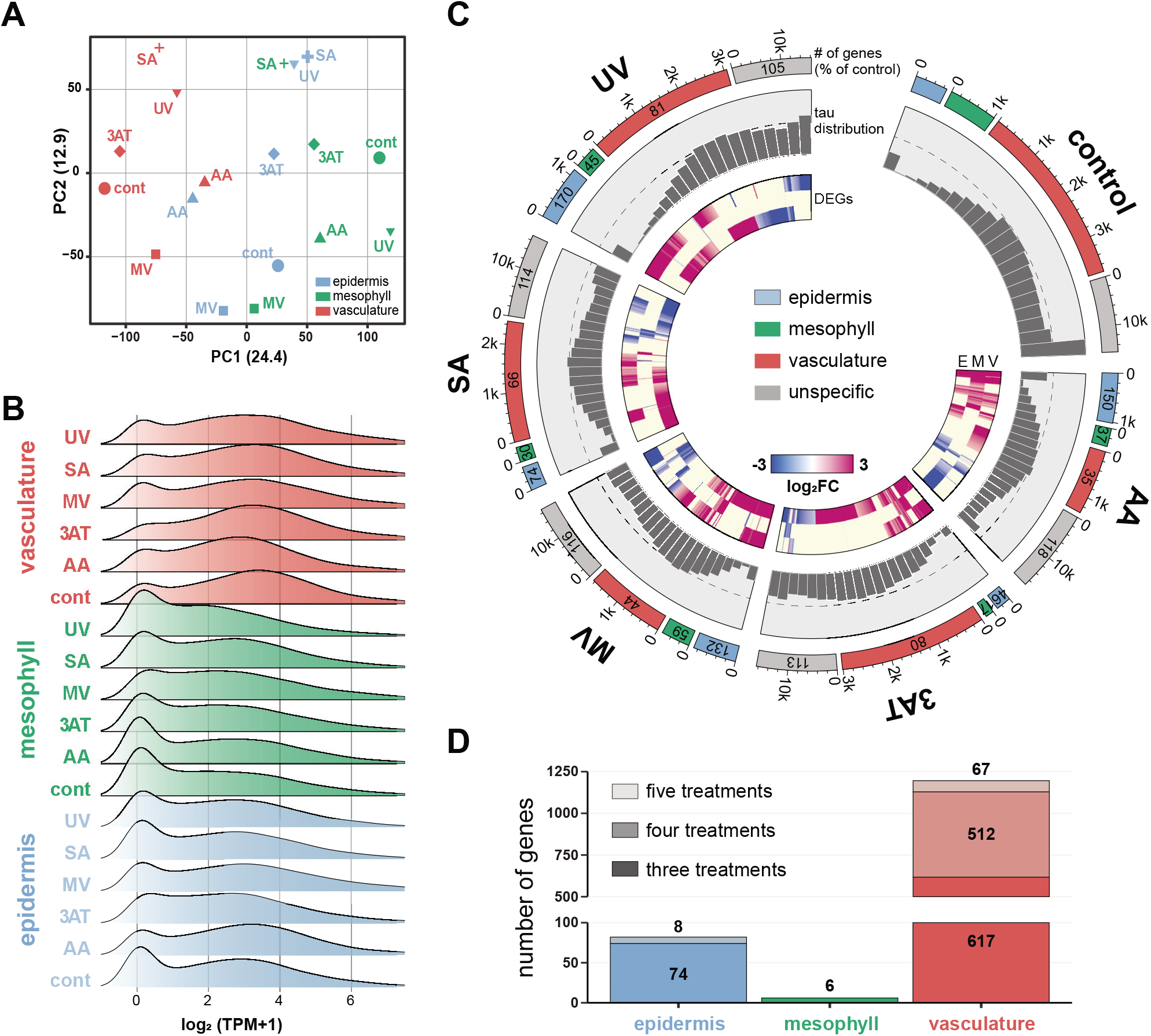
Tissue-specific expression of genes in Arabidopsis leaves after treatments with four chemical stimuli and UV-C. **(A)** Principle component analysis (PCA) for all analysed tissues and treatments. **(B)** Distribution of gene expression (in transcripts per million, TPM) in all analysed tissues after treatments. **(C)** For the individual treatments, the circular plot shows the number of genes specific for each tissue (outer scale, note the different scale for unspecific genes) and their percentage of the control (numbers in the boxes) (outer track), a histogram for the distribution of tau values of all genes (middle track; values of 0 to 1 with a bin width of 0.05) and a heatmap for the differentially expressed genes (inner track; colour scale represents the log_2_ of the fold changes). In all treatments the highest number of tissue-specific genes was in the vasculature and lowest in the mesophyll. The control had the most tissue-specific genes while after treatments the distribution of tau was more centred, i.e. the relative number of tissue-specific genes was lower. **(D)** Number of genes showing conserved tissue specificity for three, four or all five treatments, respectively. Abbreviations: AA, antimycin A; 3AT, 3-amino-1,2,4-triazole; MV, methyl viologen; SA, salicylic acid; UV, ultraviolet light.

We next asked if tissue-specific expression of genes observed under control conditions is conserved after the treatments. Compared to the control samples, the number of tissue-specific genes declined in the tissues after all five treatments between approx. 20 to 80% leading to corresponding increases in unspecific genes between 5 and 18 %. An exception was the epidermis after AA, MV and UV treatment showing increases of 50 %, 32 % and 70 %, respectively (Figure 2C, outer circles). Averaged across all tissues, the number of tissue-specific genes declined by approx. 40 % for AA, 3AT, MV and SA, and only 15 % for UV. A general decrease in tissue-specificity was also indicated by the tau value distributions which became more centred for treatments and lost the peak observable for genes with high tau values in the controls (Figure 2C, middle circle). In the vasculature 67 genes conserved their tissue-specificity across all five treatments, 579 genes after four treatments and 1,196 genes after three treatments (Figure 2D). Eight and 82 genes remained epidermis-specific after four and three treatments, respectively, while only six genes remained mesophyll-specific after three treatments. (Figure 2D). A reduction in the number of tissue-specific genes was also observed for hypoxia stress across root and shoot (Mustroph et al., 2009) and in a recent single-cell RNA-seq experiment of heat-stressed roots (Jean-Baptiste et al., 2019). In the latter a down-regulation of cell marker genes and the induction of a general stress response in all roots cells after heat shock decreased the crucial ability to assign cell identities. Although computational methods to correct for this problem are being developed (Butler et al., 2018; Jean-Baptiste et al., 2019), this remains a major challenge for the analysis of scRNA-seq experiments and highlights an advantage of laser-capture microdissection approaches for the analysis of samples from stressed plants.

Of the 14,297 genes showing no tissue-specific expression in the control samples, the vast majority remained unspecific after stress treatments, ranging from 12,289 (85 %) to 13,175 (91 %) genes for the UV treatment and the MV treatment, respectively (Figure 3A, Supplemental Table 3 and 4). The largest change from unspecific to tissue-specific expression was seen for the UV treatment for which 8 % became vasculature-specific, while the median was 3 % (Figure 3B, Supplemental Table 3 and 4). For the genes with tissue-specific expression in control conditions, a high percentage, ranging from 48 % (vasculature/UV treatment) to 82 % (mesophyll/SA treatment), changed from tissue-specific to unspecific expression, with the changes for mesophyll-specific genes significantly higher (76 %) than for the other two tissues (epidermis 59 %, vasculature 54 %) (Figure 3C, Supplemental Table 3 and 4). The remainder of tissue-specific genes stayed tissue-specific after stress treatments. For the epidermis and vasculature, the highest percentage of genes preserving their tissue specificity was on average 22 % for epidermis and 35 % for vasculature, while this was the case for only 6 % of mesophyll-specific genes (Figure 3D, Supplemental Table 3 and 4). A varying number of genes changed their tissue-specificity: ranging from 2 to 14 %, 5 to 19 % and 1 to 8 % of genes that were specific for the epidermis, mesophyll and vasculature, respectively, before stress treatments.

**Figure 3.**
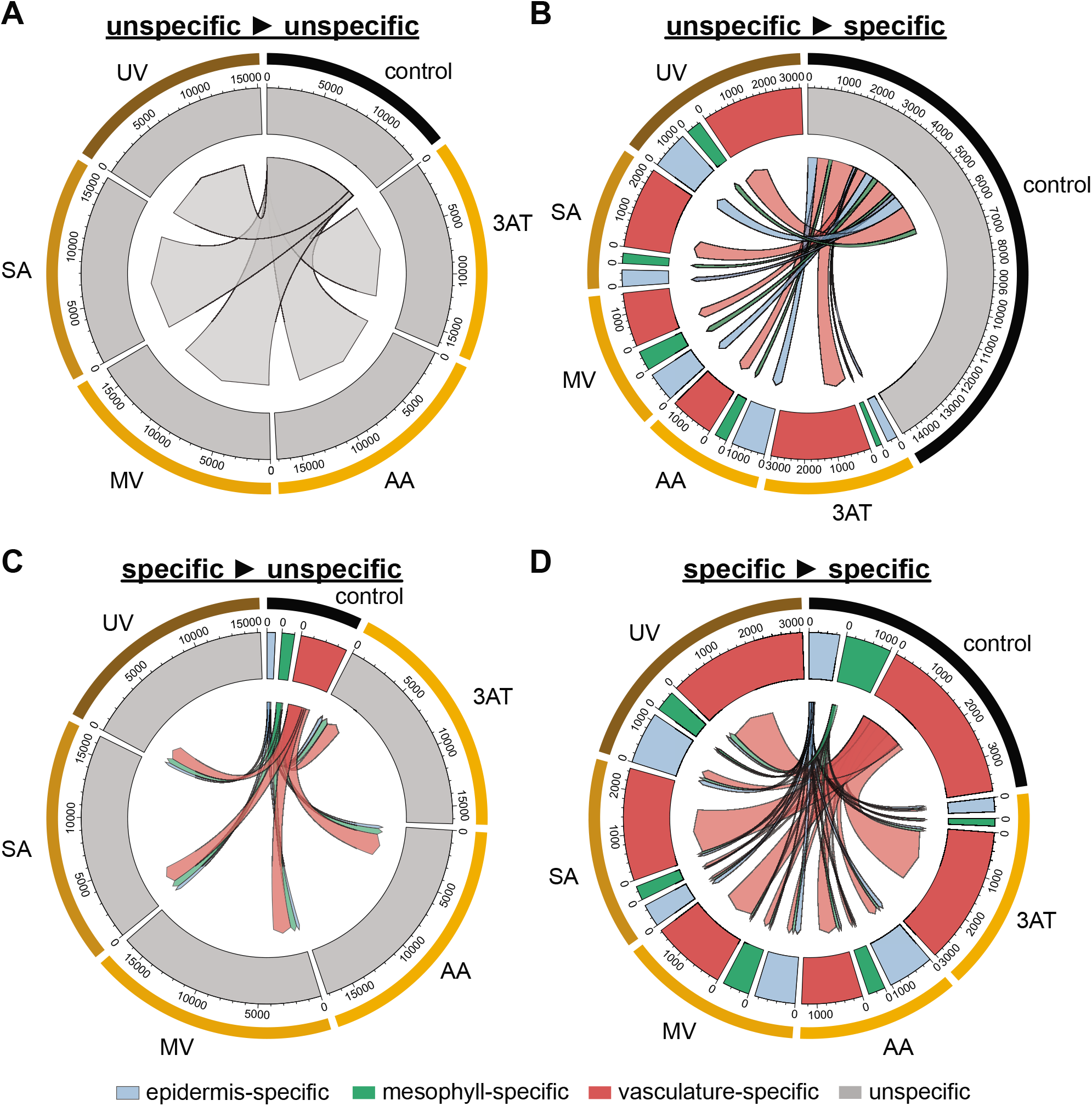
Changes in tissue-specific expression of genes after treatments with four chemical stimuli and UV-C. The tissue-specific expression of genes in control conditions was compared to treatments and results are presented as circular plots. The outer tracks indicate the six different conditions (control and five treatments) and the middle tracks the number of genes showing tissue-specific (tau≥0.85) or unspecific (tau<0.85) expression (blue: epidermis-specific; green: mesophyll-specific; red: vasculature-specific; grey: unspecific; Supplemental Table 4). The ribbons in the middle of the plots represent the number of genes and their changes in tissue-specificity after stress treatments (Note: the ribbons do not indicate gene identify information). The plots show the four possible changes in tissue-specificity: **(A)** genes remaining unspecifically expressed after stress treatment, **(B)** genes becoming tissue-specific after stress treatment, **(C)** genes loosing tissue specificity after stress treatment, and **(D)** genes that remain specific after stress treatments, with some changing their tissue-specificity. Abbreviations: AA, antimycin A; 3AT, 3-amino-1,2,4-triazole; MV, methyl viologen; SA, salicylic acid; UV, ultraviolet light.

Our LCM-seq data provide robust marker genes conserving their tissue-specificity after diverse treatments, which is important for the determination of cell identities in future scRNA-seq experiments. The results also show a general decrease in tissue-specificity of genes after stress treatments. This indicates that cells switch from performing their specific function within an organ to a more generalised survival response, in agreement with earlier findings (Jean-Baptiste et al., 2019; Mustroph et al., 2009).

### Differential stress responses of genes across tissues

After determining the changes in tissue specific expression of genes in response to treatments, we next analysed quantitative changes in gene expression levels irrespective of tissue specificity. The number of differentially expressed genes (DEGs; |log2 (fold change)| >1, false discovery rate (FDR) < 0.05) for each stress treatment varied across the tissues when compared to the controls. In total 3,793 DEGs were identified across all treatments in the three tissues (Supplemental Table 5), with the epidermis generally showing the highest number of DEGs, while the mesophyll had the lowest number (Figures 4A). For antimycin A the responses were similar across tissues as indicated by a comparable number of DEGs (mesophyll: 579 DEGs, vasculature: 521 DEGs, epidermis: 389 DEGs), while UV irradiation had the largest effect on the epidermis (1,979 DEGs) and vasculature (1,541 DEGs) and its effect on the mesophyll was limited (15 DEGs). It is possible that the mesophyll is better protected from UV stress by nuclear photo-relocation to avoid DNA damage. The movement of the nucleus to the side-wall of mesophyll cells reduces exposure to UV light and this is accentuated by shielding of the nucleus by chloroplasts (Iwabuchi et al., 2016; Suetsugu et al., 2015). Avoidance movement of chloroplasts, mainly observed for excessive high- and blue light (Kasahara et al., 2002), but also after UV stress (Hermanowicz et al., 2019), might also reduce the production of ROS and hence explain the low number of DEGs in the mesophyll observed after UV irradiation. In the epidermis, DEGs overlapping for at least two out of the five treatments made up the four largest intersects of the DEG lists (Figure 4A). By contrast, DEGs were markedly more treatment-specific in the other two tissues, with the three and four largest intersects only contained DEGs responding to a single treatment in the mesophyll and vasculature, respectively (Figure 4A). DEGs shared across all treatment were only present in the vasculature (136 DEGs). Thus, although some genes respond to two or more treatments in the different tissues, there is no evidence for a significant, common stress response in any given tissue. Similar results were obtained in Arabidopsis root tissues after several stress treatments (Iyer-Pascuzzi et al., 2011).

**Figure 4:**
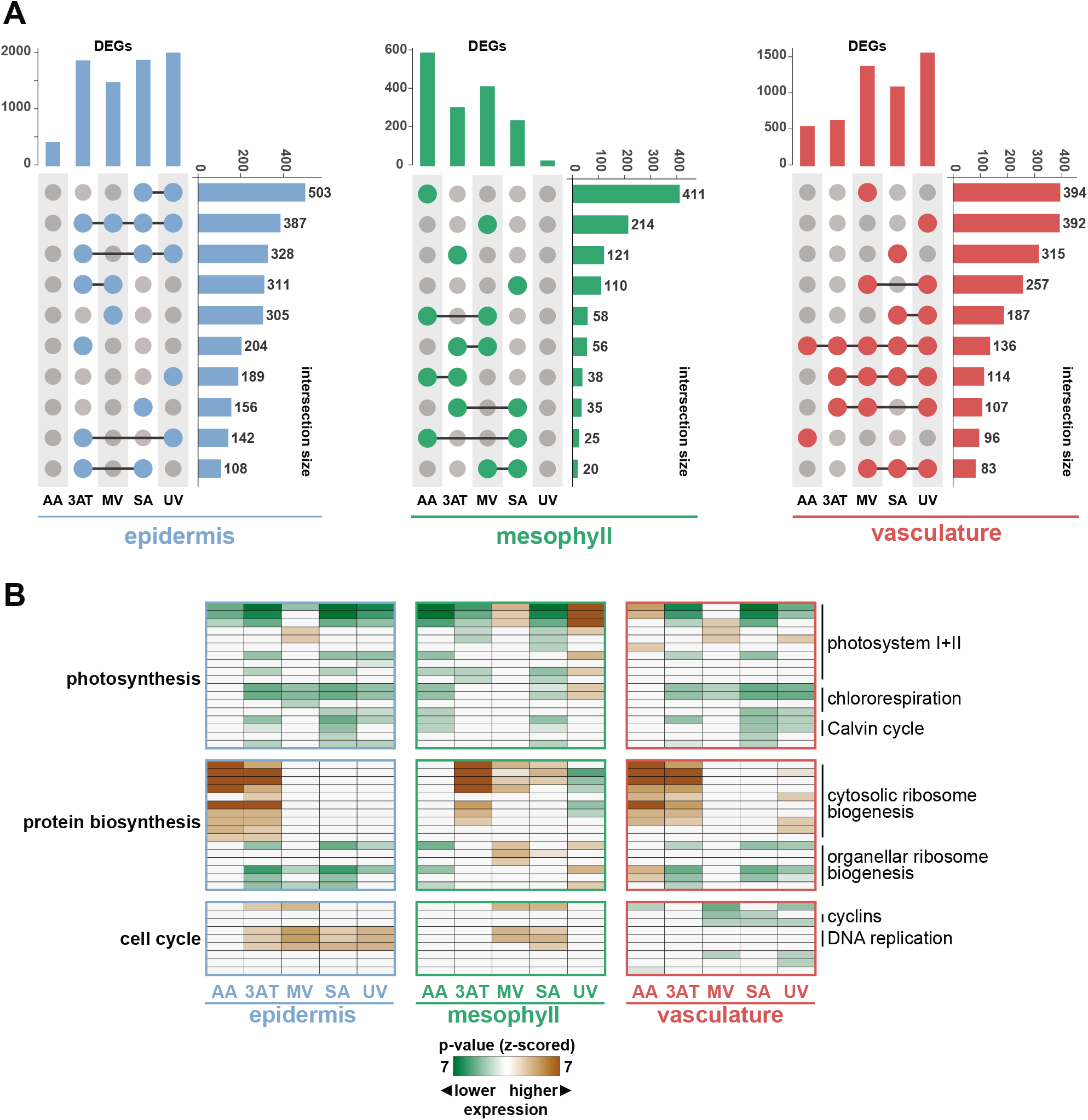
Overlaps in differentially expressed genes and their overrepresentation in functional categories. (A) The numbers and overlaps of differentially expressed genes (DEGs; |log_2_ (fold change)| > 1, FDR < 0.05) were determined and are shown as UpSet plot representations (Conway et al., 2017). In each of the plots the vertical bar chart gives the number of DEGs when compared to the untreated controls, while the number of overlapping DEGs across treatments is represented by horizontal bars, with the dot matrix indicating the respective overlaps by connected circles. For example, for the epidermis the comparison against the control showed that 503 DEGs overlapped for the SA and UV treatments, while 305 DEGs were specific for the MV treatment. Shown are only the 10 largest intersects. (B) Using the PageMan tool, the difference in expression of genes in functional categories was determined by comparison of the stress treatments with the control and is represented by heatmaps (Usadel et al., 2006). Indicated on the left are the annotations of high order bincodes and on the right selected, more detailed bincodes (see Supplemental Table 6 for complete listing). The colour scale indicates the p-value for statistical difference (Wilcox rank-sum test, z-scored after Bonferroni correction) of the median fold-change of genes in the individual bins to all other genes. Brown and green colour indicate higher and lower median expression, respectively, and thus stronger stress responses of genes in these functional bins. Abbreviations: AA, antimycin A; 3AT, 3-amino-1,2,4-triazole; MV, methyl viologen; SA, salicylic acid; UV, ultraviolet light.

To further probe for similar and contrasting responses of the tissues to treatments, the PageMan tool with a Wilcox rank sum test was applied on the full data set (Usadel et al., 2006). This analysis identified functional categories for which the corresponding genes show statistically significant (p < 0.05 after Bonferroni correction) higher up- or down-regulation than all other genes. This revealed complex and distinct response patterns of tissues to the five treatments, with the full results of this analysis shown in Supplemental Figure 2 and listed in Supplemental Table 6. Here we highlight only three fundamental functional categories (‘photosynthesis’, ‘protein biosynthesis’, ‘cell cycle’) (Figure 4B). Genes in the category ‘photosynthesis’ (including photosystem I and II, chlororespiration and calvin cycle) were downregulated in all three tissues after 3AT and SA treatments, while photosystem I and II-related genes were up-regulated by MV in accordance with its inhibitory function on photosystem I. UV irradiation generally led to a down-regulation of photosynthesis-related genes in epidermis and vasculature, but a strong up-regulation in the mesophyll (Figure 4B). This was somewhat surprising given the low number of DEGs in the mesophyll for this treatment (Figure 4A). The basis for this result is the concerted, but minor, up-regulation of many genes in this functional category not classified as DEGs by our stringent cut-off (|log2 (fold change)| > 1, FDR < 0.05), leading to a high statistical significance and highlighting the value of this categorical, global analysis (Usadel et al., 2006). A contrasting response was also observed for AA, with a down-regulation of genes in this category for the epidermis and especially the mesophyll, and up-regulation in the vasculature. The transcriptional responses of the epidermis and the vasculature for the cytosolic ribosome biogenesis-related genes was very similar, with high expression after AA and 3AT. In the mesophyll there was also higher gene expression for 3AT but not for AA as in the other two tissues. In addition, there was higher expression of these genes in response to MV and SA, while lower for UV. For the latter however the expression was higher in the vasculature. The expression of genes associated with organellar ribosome biogenesis was higher than other genes in the epidermis and vasculature after 3AT, SA and UV treatments, and lower in the mesophyll for MV and UV. For genes involved in the cell cycle there was higher expression of these genes in the epidermis after 3AT, UV, SA and MV and for the latter two also in the mesophyll, while their expression was lower in all treatments except 3AT in the vasculature.

Together these results indicate that the stress treatments led to distinct changes in expression patterns in the leaf tissues. The epidermis and the vasculature generally showed more similar responses, i.e. up- or down-regulation of genes in the same functional categories under the same stresses, and these were often contrasting with the mesophyll. However, there was no evidence for an extensive set of common stress-responsive genes across all tissues. Together, this highlights how the analysis of whole tissue samples might be a read-out of a strong response in only one tissue or a mixed read-out of specific responses of multiple tissues.

### Tissues show common and specific responses to AA treatment

We next analysed the extent of specific and overlapping responses of tissues to the individual treatments. The epidermis had the highest number of specific DEGs for 3AT, SA and UV, and the epidermis and vasculature shared a significant number of DEGs that made up between 21 % (epidermis/3AT) to 66 % (vasculature/SA) of all their DEGs (Figure 5A). Therefore, and in contrast to limited conservation of stress-responses across treatments in a given tissue (Figure 4A), for any specific treatment there was some conservation of DEGs across tissues, especially for epidermis and vasculature (Figure 5A).

**Figure 5:**
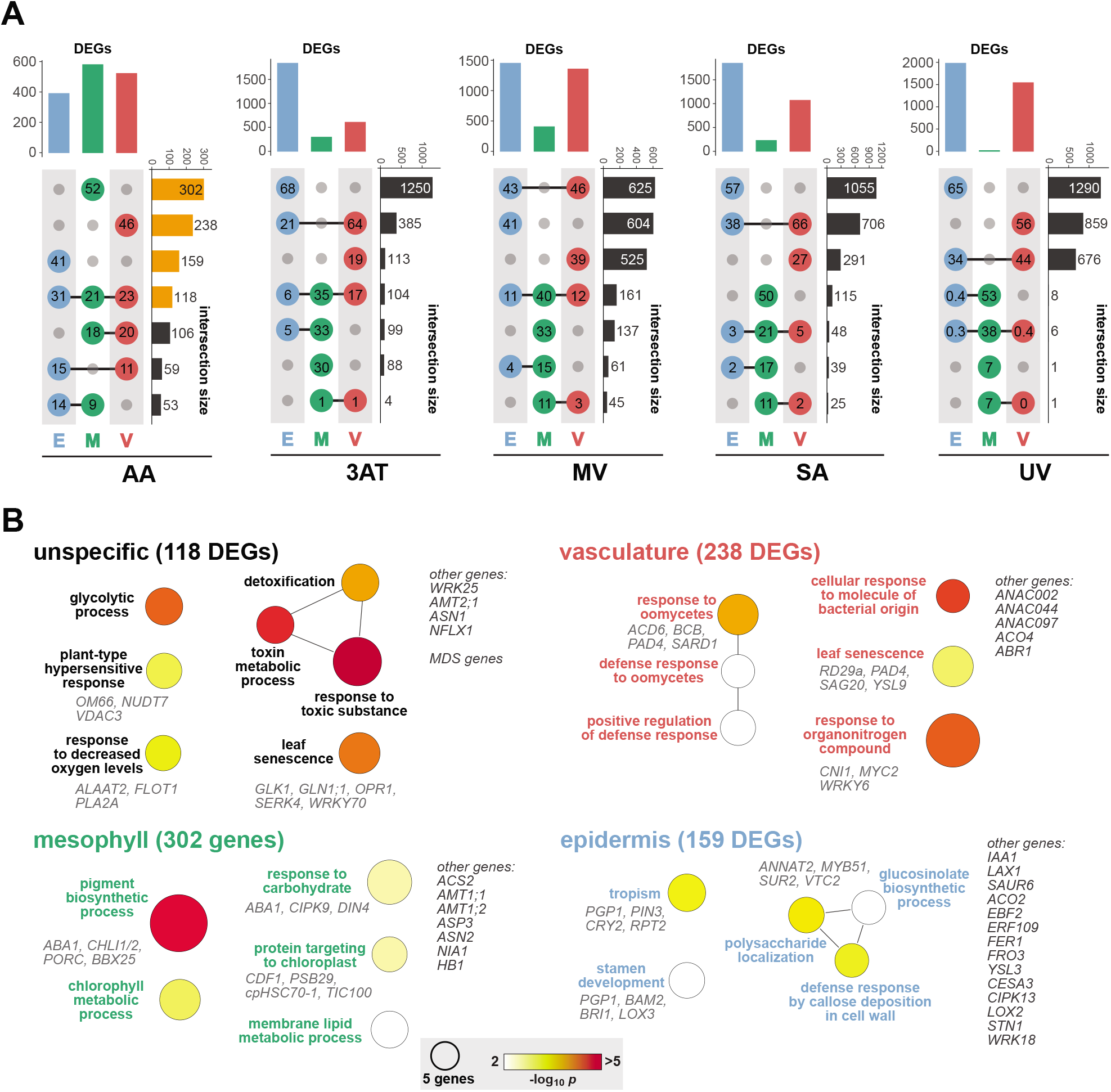
Differential responses in gene expression in leaf tissues after treatments with four chemical stimuli and UV-C. (A) Upset plot representation for overlapping sets of DEGs in three analysed tissues for each of the five treatments. In each plot the vertical bar chart gives the number of DEGs in the three tissues (E, epidermis - blue; M, mesophyll - green; V, vasculature - red) while the number of overlapping DEGs across tissues is represented by the horizontal bar chart, with the dot matrix indicating the respective intersect by connected circles. The numbers in the dots give the percentage of DEGs in the given intersect. For example, after antimycin A (AA) treatment 302 DEGs were specific for the mesophyll and 119 DEGs were shared in all three tissues. Vertical bars highlighted in orange for the AA treatment were further analysed for their GO term enrichment. (B) GO term enrichment analyses for the DEGs common between, or specific for, each of the three tissues, respectively, after AA treatment (indicated by vertical bars in orange colour in A). The size of circles indicates the number of genes included in each enriched (p<0.01) GO term and the colour scale the corresponding p-value after Bonferroni correction. See Supplemental Table 6 for details. Abbreviations: AA, antimycin A; 3AT, 3-amino-1,2,4-triazole; MV, methyl viologen; SA, salicylic acid; UV, ultraviolet light.

The range of specificity and overlap of DEGs in all three tissues after AA treatment (Figure 5A), in conjunction with the contrasting responses of functional categories as described above (Figure 4B), made this treatment a focus for a more detailed and comprehensive analysis. This also provides insight into the yet unexplored tissue-specific transcriptional responses to mitochondrial dysfunction. The largest intersects were those containing DEGs specific to each of the three tissues (52 %, 46 % and 41 % of DEGs specific to the mesophyll, vasculature and epidermis, respectively; Supplemental Table 7) followed by the intersects containing the 118 DEGs shared amongst all (21 %, 23 % and 31 % of DEGs in mesophyll, vasculature and epidermis; Supplemental Table 7) (Figure 5A, bars highlighted in orange).

For these 118 DEGs enriched GO terms were related to glycolysis, leaf senescence, toxin metabolism and hypersensitive response (Figure 5B), which, together with the associated DEGs, are generally established on a whole leaf tissue to represent the impact of AA treatment (Ivanova et al., 2014; Ng et al., 2013). A set of marker genes highly induced by AA are the mitochondrial dysfunction stimulon (MDS) genes which are controlled by the key regulators of mitochondrial retrograde signalling ANAC013 and ANAC017 (De Clercq et al., 2013; Ng et al., 2013). With the exceptions RPL12 and AT5G14730, the MDS genes detectable in our RNA-seq data showed significant increases of their transcript abundances in the response to AA in all tissues (Figure 6). While some MDS genes had similar transcript abundances in control conditions, e.g. AOX1a, At12CYS2 and HRE2, other genes showed significant variation under control conditions. For example, ANAC013 and OUTER MITOCHONDRIAL MEMBRANE PROTEIN OF 66 KDA (OM66) showed an at least 10-fold difference in expression between the vasculature and mesophyll, while UP-REGULATED BY OXIDATIVE STRESS (AT2G21640) was not expressed in the epidermis and highest in the vasculature. The induction of MDS genes after AA treatment differed across the tissue for some of the MDS genes. The fold changes for AOX1a, At12CYS-2 and UGT74E2 were similar in all three tissues, whereas for genes such as UPOX, HSP23.5 or ABCB4 the differences varied by up to 20-fold (Figure 6).

**Figure 6:**
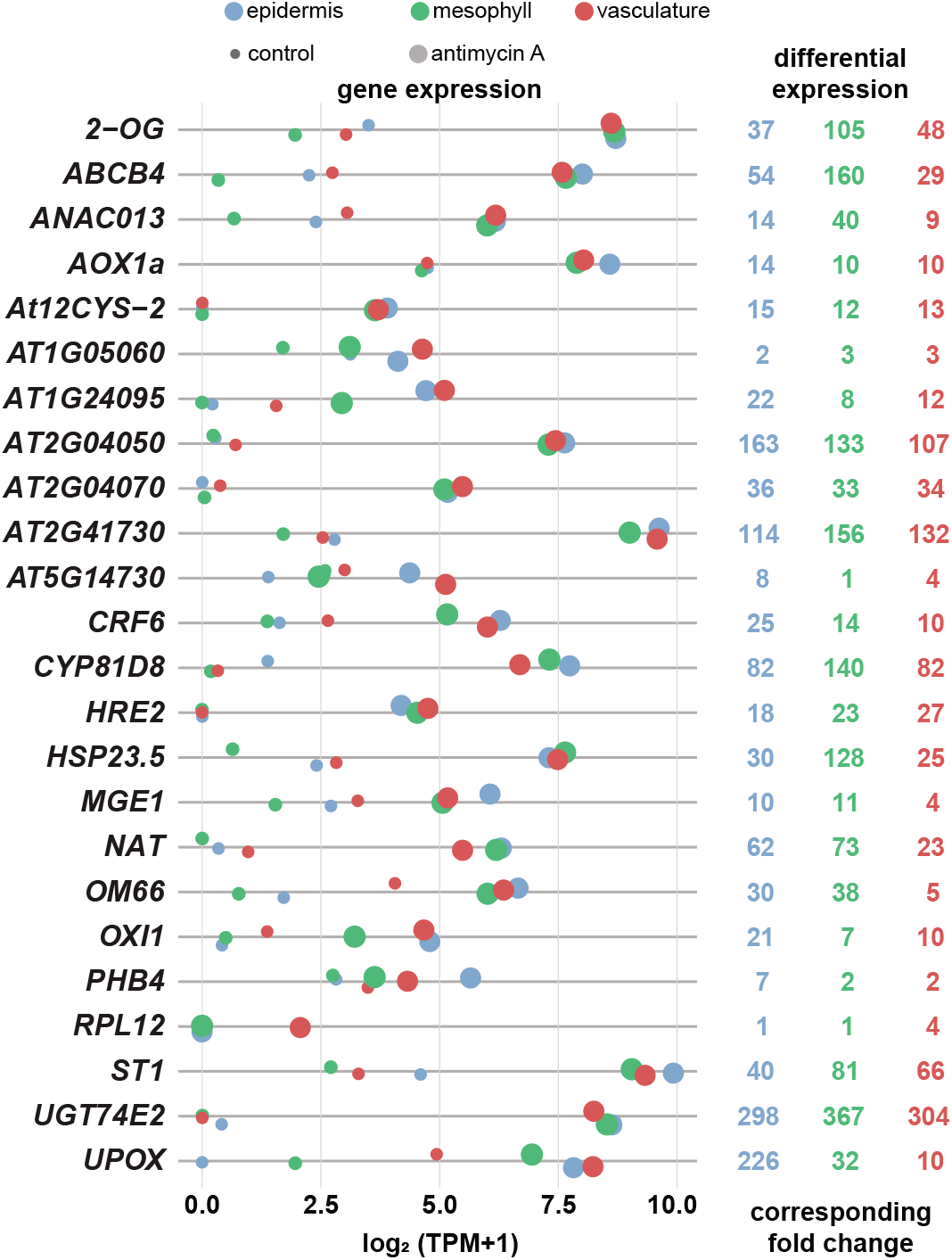
Increased expression of mitochondrial dysfunction stimulon genes in three leaf tissues after antimycin A treatment. The expression of genes known to respond to mitochondrial dysfunction (mitochondrial dysfunction stimulon, MDS; De Clercq et al 2013) is given in the dot plot for the three analysed leaf tissues under control conditions and after antimycin A (AA) treatment. Expression is given based on transcripts per million (TPM) as log_2_(TPM+1). The values on the right give the fold induction of each gene in the three tissues after AA treatment.

For the 302 genes differentially expressed after AA treatment in the mesophyll only (Figure 5A), a GO term enrichment analysis identified terms and genes related to a down-regulation of chlorophyll, lipid and carbohydrate metabolism as well as chloroplast protein import (Figure 5B, Supplemental Table 8). The list of DEGs also included a number of genes with a function in nitrogen assimilation and metabolism: *AMMONIUM TRANSPORTER 1;1 (AMT1;1*, 2.5-fold up-regulated), *AMT1;2* (30-fold down-regulated), *ASPARTATE AMINOTRANSFERASE 3* (*ASP3*, 2.7-fold down-regulated), *ASPARAGINE SYNTHETASE 2* (*ASN2*, 3-fold down-regulated) and *NITRATE REDUCTASE 1* (*NIA1*, 4.5-fold down-regulated) (Supplemental Table 5). This general downregulation of metabolic pathways, also evident from the PageMan analyses (Figure 4B, Supplemental Figure 2), indicates that mitochondrial dysfunction induced by AA impacts the assimilation of nitrogen and carbohydrates into amino acids and downstream metabolites such as pigments in the mesophyll. Among the mesophyll-specific DEGs most-highly up-regulated by AA were *HEMOGLOBIN 1* (*HB1*, 100-fold up-regulated) and *1-AMINOCYCLOPROPANE-1-CARBOXYLIC ACID SYNTHASE 6* (ACS6, 30-fold up-regulated) (Supplemental Table 5). Both genes are also highly up-regulated on a whole tissue level by ethylene, hypoxia and submergence (Hartman et al., 2019; Meng et al., 2020; Peng et al., 2005; Trevaskis et al., 1997), in line with evidence that the transcriptional responses to AA and hypoxia are overlapping (Wagner et al., 2018). This suggests that AA induces stresses related to local changes in oxygen levels specifically in the mesophyll, while AA has also a putative role in impeding photosynthetic efficiency by inhibiting cyclic electron flow (Labs et al., 2016). These results are in agreement with recent findings showing ethylene is mediating hypoxia tolerance via nitric oxide (NO) (Hartman et al., 2019). In accordance with this, the downregulation of NIA1 and up-regulation of HB1 by ethylene, produced by increased ACS6, would decrease NO levels and thus inhibit the degradation of the oxygen-sensing group VII ERFs through the N-end rule pathway. These ERFs are the major transcription factors activating the expression of hypoxia-related genes (van Dongen and Licausi, 2015). One of these, HYPOXIA RESPONSIVE ERF 2 (HRE2), is a member of the highly induced mitochondrial dysfunction stimulon genes and its expression is directly controlled by the master regulators of mitochondrial retrograde signalling ANAC013 (De Clercq et al., 2013) and ANAC017 (Meng et al., 2019; Ng et al., 2013). Our LCM-seq data shows a 30-fold up-regulation of HRE2 in all tissues after AA treatment (Figure 6), similar to a its observed upregulation in different cell types under hypoxia (Mustroph et al., 2009). This presents a direct link for the co-expression of genes involved in mitochondrial stress and hypoxia mediated by ethylene.

GO terms enriched for the 238 vasculature-specific DEGs (Figure 5A) were related to defence responses, leaf senescence and nutrient balancing (Figure 5B). The latter included *CARBON/NITROGEN INSENSITIVE 1* (*CNI1*) and *WRKY6*. The up-regulation (above 16-fold) of these two genes suggests a change in nutrient balance within the vasculature after AA treatment as CNI1 is involved in maintaining the C/N balance (Sato et al., 2009) and WRKY6 in phosphate homeostasis (Ye et al., 2018). Similar to the connection of AA responses with hypoxia- and ethylene-related signalling seen for the mesophyll, the genes encoding the ethylene biosynthetic enzyme 1-AMINOCYCLOPROPANE-1-CARBOXYLATE OXIDASE 4 (ACO4) and the ERF transcription factor ABSCISIC ACID REPRESSOR1 (ABR1), up-regulated 2- and 30-fold (Supplemental Table 5), respectively, are also among the vasculature-specific DEGs. ABR1 promotes auxin biosynthesis (Ye et al., 2020) and ethylene-induced auxin biosynthesis provides a means for hormonal crosstalk between ethylene and auxin (Stepanova et al., 2005).

For the 159 AA-responsive DEGs specific to the epidermis, enriched GO terms were ‘tropism’, ‘stamen development’ and ‘polysaccharide localisation’ (Figure 5B). Among these DEGs were down-regulated genes *PIN-FORMED3* (*PIN3*) and *LIKE AUXIN RESISTANT 1* (*LAX1*), both encoding auxin transporters (Friml et al., 2002; Peret et al., 2012), as well as up-regulated *SUPERROOT2* (*SUR2*) involved in auxin homeostasis (Barlier et al., 2000) and *SMALL AUXIN UPREGULATED RNA 6* (*SAUR6*) down-regulated 90-fold (Supplemental Table 5). Moreover, the *INDOLE-3-ACETIC ACID INDUCIBLE 1* (*IAA1*) gene, encoding a negative regulator of auxin responses from the AUX/IAA protein family (Yang et al., 2004), was down-regulated about 30-fold (Supplemental Table 5). The transcript abundances of *PIN3, LAX1* and *IAA1* are controlled by auxin levels (Paponov et al., 2008), and hence their down-regulation after AA treatment, together with the auxin-induced marker *SAUR6*, suggests decreased auxin levels in the epidermis. Together with a reduced transport of auxin into the epidermis under stress indicated by down-regulation of *PIN3* and *LAX1*, this would remove inhibition of retrograde signalling to increase expression of mitochondrial stress-responsive genes (Ivanova et al., 2014; Kerchev et al., 2014). Another PIN protein, PIN1, was already reported previously as a regulator of retrograde signalling (Ivanova et al., 2014). In addition to auxin, DEGs associated to ethylene synthesis (*ACC OXIDASE 2*, *ACO2*) and transcriptional regulation (*EIN3-BINDING F BOX PROTEIN 2, EBF2; ETHYLENE RESPONSE FACTOR 109, ERF109; RELATED TO AP2 4, RAP2.4*) were also in this gene list. *ERF109, ACO2* and *RAP2.4* were down-regulated by 30-fold, 3-fold and 2-fold, respectively, while *EBF2* was up-regulated by 3-fold (Supplemental Table 5). The overexpression of ERF109, also termed REDOX RESPONSIVE TRANSCRIPTION FACTOR 1 (RRTF1) due to its initial identification as regulator of redox homeostasis (Khandelwal et al., 2008), leads to auxin over-production (Cai et al., 2014). Hence its strong down-regulation (30-fold) is also indicative for a reduction in auxin levels under AA treatment and decreased ethylene production by downregulation of ACO2 would also reduce auxin uptake into the epidermis (Lewis et al., 2011; Vandenbussche et al., 2010). Furthermore, increased expression of EBF2 in the epidermis has been shown to decrease the expression of the auxin importer AUXIN RESISTANT 1 and restrict plant growth via the epidermis (Vaseva et al., 2018).

Taken together, these results provide novel insights into tissue-specific stress responses. Specifically for the AA treatment, while MDS genes play an important role in mitigating mitochondrial dysfunction and other stresses such as hypoxia in all tissues (Wagner et al., 2018), they contribute to varying degrees in different tissues and also show variation of their expression levels under normal growth conditions. We provide first evidence that the interaction of auxin and ethylene mediates retrograde signalling during AA-induced mitochondrial dysfunction, which has not been observed before on a whole tissue level.

### Tissue-Specific and Unspecific Gene Regulatory Networks

The tissue-specific responses of genes to AA treatment, together with differences in the magnitude of response of the MDS genes, suggested a tissue-specific regulation of gene expression. We created a gene regulatory network (GRN) to identify putative underlying transcriptional networks and their associated regulatory TFs (regulators) using the TF2network tool (Kulkarni et al., 2018). The GRN was generated by applying robust enrichment statistics (FDR < 0.01) on experimentally confirmed TF binding sites and their occurrence in the tissue-specific and unspecific DEGs identified for the AA treatment. This identified 23 GRN regulators with the underlying network containing 252 nodes and 922 edges with an average number of neighbours of 7.3 (Supplemental Table 9). The GRN was visualised by applying a semi-hierarchical, radial layout (Figure 7). For the mesophyll-specific DEGs no upstream regulators were enriched suggesting these genes were not controlled by a set of common TFs or by TFs with unknown binding motifs. This may suggest that the response in mesophyll cells is a secondary response due to inhibition of mitochondria resulting in inhibition of chloroplast metabolism and function as evidenced by the GO enrichment analyses (Figure 5).

**Figure 7:**
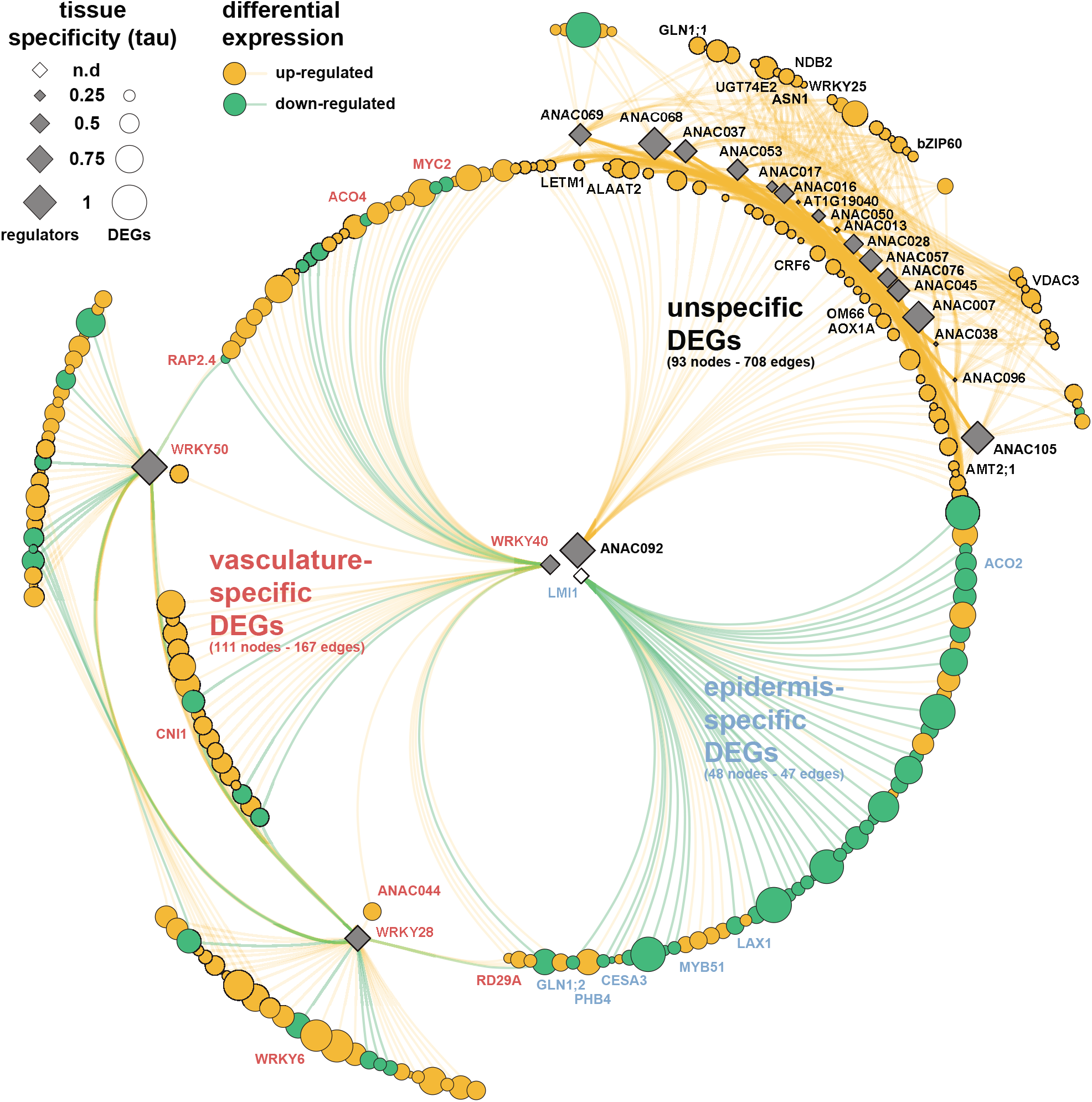
Gene regulatory network for differentially expressed genes in the three analyses tissues after antimycin A treatment. For genes differentially expressed in the epidermis (blue annotations), vasculature (red annotations) or unspecifically in all tissues (black annotations) a gene regulatory network (GRN) was generated using the T2FNetwork tool (Kulkarni et al., 2018). For the 302 mesophyll-specific DEGs no regulators were identified by this method. Out of the remaining 494 DEGs, 21 upstream transcription factors were determined as candidate upstream regulators (diamonds) for 231 DEGs (circles; up-regulated by AA treatment in orange, down-regulated in green). The GRN is presented as a semi-hierarchical radial layout of the 252 nodes and 922 edges using the yfiles algorithm and edge-bundling. For each gene the sizes of circles and diamonds represent their tissue-specificity (i.e. tau values) in the AA treatment. Names of genes discussed in the text are indicated.

For the unspecific DEGs their 18 predicted up-stream regulators were exclusively ANAC transcription factors, with ANAC013 being the only regulator differentially expressed itself after AA treatment (between 9- and 40-fold; Figure 6). Most DEGs had several ANACs as their regulators in highly combinatorial relationships (Figure 7). These ANACs included 4 out of the 5 ER-tethered ANACs known to bind the mitochondrial dysfunction motif (ANAC013, ANAC016, ANAC017, ANAC053) (De Clercq et al., 2013) and in agreement with their function as transcriptional regulators were connected to up-regulated DEGs. All also had low tissue-specific expression with *ANAC013* having the lowest (tau = 0.05) and *ANAC053* the highest (tau = 0.54) among them (Figure 7), indicating a generalised function of these ANACs in the response to AA across tissues. By contrast, other ANACs such as *ANAC092* and *ANAC105*, had an almost exclusive expression (tau = 0.99 and tau = 0.91, respectively) in one tissue, suggesting they modulate the induction of these DEGs driven by other ANACs with lower tissue specificity such as *ANAC013* or *ANAC017*. While the ANACs were interconnected among themselves and the DEGs, ANAC092 shared edges with a subset of DEGs, among them AA-response marker genes *AOX1a* and *OM66*. ANAC092 is a positive regulator of leaf senescence (Kim et al., 2009) and highly up-regulated in ANAC017 overexpressing lines showing early leaf senescence and cell death (Meng et al., 2019). It is also a target of EIN2 and EIN3 in the ethylene response (Chang et al., 2013; Kim et al., 2014). Our GRN suggests that ANAC092 may control the expression of a subset of AA responsive DEGs on a tissue-specific level mediated by ethylene.

For the epidermis-specific DEGs only the homeodomain TF LATE MERISTEM IDENTITY1 (LMI1) was a predicted as up-stream regulator (Figure 7). The *LMI1* gene showed very low expression levels in our LCM-seq data and hence its tissue specificity and differential expression could not be determined reliably. This is in agreement with its function in leaf shape determination by repressing local growth during early meristem differentiation (Vlad et al., 2014). Putative target genes of LMI1 include *ACO2* and *LAX1* indicating a role of ethylene and auxin for its function. Another LMI1 target gene up-regulated by AA treatment in the epidermis encoded PROHIBITIN 4 (PHB4), a member of the prohibitin family of mitochondrial proteins and also part of the MDS genes (Figure 6). These proteins are important for meristem function and leaf development and knock-out of a closely related prohibitin, PHB3, accentuates retrograde signalling (Van Aken et al., 2010). Therefore, LMI1 could control developmental aspects of the response to mitochondrial dysfunction induced by AA treatment in the epidermis. A constitutive up-regulation of mitochondrial retrograde signalling, for example by overexpression of ANAC017, leads to growth retardation and early senescence (Meng et al., 2019), highlighting the link between mitochondrial signalling and plant development.

Three regulators were identified for the vasculature-specific DEGs in the GRN: WKRY28, WRKY40 and WRKY50 (Figure 7). In contrast to the highly interconnected tissue-unspecific network (average neighbour number = 15), the three WRKYs were mainly up-stream regulators of DEG subsets with limited interconnectivity (average neighbour number = 3). WRKY40 is a repressor of mitochondrial signalling and is also involved in the co-ordination with chloroplast signalling (Van Aken et al., 2013). In addition, WRKY40 has a function in the regulation of touch and submergence responses, with both stress pathways in part mediated by ethylene (Meng et al., 2020; Xu et al., 2019). Correspondingly, among the down-regulated DEGs potentially controlled by WRKY40 in the vasculature were the ethylene biosynthesis enzyme-encoding *ACO4* (see above) and the transcription factors RAP2.4 and MYC2. RAP2.4 is involved in the jasmonic acid (JA) and ethylene-mediated defence responses (Sham et al., 2019). Similarly, MYC2 is also involved in the regulation of jasmonic acid/ethylene cross-talk by directly interacting with the master regulators of ethylene signalling, EIN3 and EIN3-LIKE1 (Song et al., 2014; Zhang et al., 2014). The GRN further suggests that WRKY40 and WRKY28 share a regulator function for ANAC044 (Figure 7). Up-regulation of ANAC044, together with ANAC085, leads to stress-induced cell cycle arrest (Takahashi et al., 2019). Given that mitosis and cell-cycle-related functional categories were highly enriched for genes with vasculature-specific expression (Supplemental Figure 1), it seems that mitochondrial dysfunction induced by AA treatment impacts growth by inhibiting cell division especially in this tissue. WRKY28 is also a putative upstream regulator of WRKY6, with the latter regulating phosphate uptake into the xylem by controlling the expression of the phosphate translocator PHOSPHATE1 (Chen et al., 2009). Thus, WRKY28 might be a regulator co-ordinating cell proliferation and nutrient homeostasis under stress conditions, and our LCM-seq data suggests this role in the vasculature. Another subset of vasculature-specific DEGs had WRKY50 as an upstream regulator in the GRN (Figure 7). WRKY50 is generally considered a transcription factor with a role in defence responses (Gao et al., 2011), but evidence also suggests it is a target of the transcription factor RESPONSE REGULATOR 2 (ARR2) (Hass et al., 2004). ARR2 is a component of the ethylene signalling pathway downstream of ETHYLENE RESPONSE 1 (ETR1) (Hass et al., 2004) and also activates the transcription of at least three components of complex I in the mitochondrial respiratory chain (55 kDa subunit NADH-binding protein, At5g08530; NADH-ubiquinone oxidoreductase 20 kDa subunit, At5g11770; NADH dehydrogenase (ubiquinone) Fe-S Protein, At1g79010) (Lohrmann et al., 2001). Thus, the GRN regulator WRKY50 might integrate diverse stress signals, including retrograde signals resulting from mitochondrial stress after the AA treatment.

Taken together, by using the response to AA treatment as a case study we generated a GRN for stress-responsive, tissue-specific genes. This approach has potential to allow for a more specific engineering of stress tolerant plants by targeting only subsets of genes and in specific tissues by employing their upstream regulators.

## DISCUSSION

The complex nature of tissue-specific responses cannot be predicted from whole organ studies and their analysis provides potential targets for optimised engineering of more stress tolerant plants (Long, 2011; McFaline-Figueroa et al., 2020). Studies analysing transcriptional responses on a tissue or cell-specific level have largely focussed on the Arabidopsis root because of their modular anatomy, linear developmental progression and methodological accessibility. Detailed comparative studies in leaf tissues using a variety of stresses are not available, although leaves are the energy and carbon capturing organs and directly affected by many abiotic and biotic stresses. The limited number of studies investigating tissue-specific aspects of stress responses of Arabidopsis roots indicate cell-type specific expression decreases under adverse conditions such as hypoxia (Mustroph et al., 2009) and heat stress (Jean-Baptiste et al., 2019). Furthermore, differentially expressed genes across stress treatments are not conserved in root cell types (Iyer-Pascuzzi et al., 2011). Similarly, we observed a reduction in the number of tissue-specific genes, i.e. a more uniform and less diverse expression of genes, in leaf tissues after applying four chemical stimuli and UV stress. For any given leaf tissue, the treatments led to significant induction or repression of genes, but these were limited to only one or few treatments and a common response was largely absent. A limited number of genes also retained, gained or changed their tissue specificity. By contrast, for each individual treatment a significant overlap of DEGs among the three tissues was observable, especially for epidermis and vasculature.

A the role of chloroplasts and mitochondria in sensing and signalling stress responses has emerged in the last decade (Crawford et al., 2017; Kleine and Leister, 2016; Wang et al., 2020). Recently evidence has emerged that the epidermis and vascular parenchyma contain sensory plastids that are smaller in size than the mesophyll chloroplasts. These epidermal plastids have a function in the response to osmotic stress and shade avoidance through auxin-regulated growth in the stem (Procko et al., 2016; Veley et al., 2012), while sensory plastids in the vasculature have been associated to ABA biosynthesis and drought stress (Endo et al., 2008; Galvez-Valdivieso et al., 2009). Concomitantly, proteomic analysis of sensory plastids suggests they integrate environmental conditions leading to changes in gene expression and epigenetic alterations (Beltrán et al., 2018). Similar to these findings, we observed a limited response of the mesophyll to the all treatments except AA, and significant changes in gene expression in the epidermis and vasculature after all treatments that might at least partly be driven by such sensory plastids.

Our data identified tissue-specific responses to mitochondrial dysfunction induced by AA not previously observed on a whole organ level. Difference in mitochondrial morphology, number and function across different cell types have been described, (Logan, 2006). In Arabidopsis, total mitochondrial number and volume increases in epidermal cells exposed to cold, whilst cristae to matrix ratio increases in the mesophyll (Armstrong et al., 2006). The number of mitochondria declines faster in epidermal than mesophyll cells under dark induced senescence (Keech et al., 2007). Although these results suggest tissue-specific mitochondrial responses and regulation, so far this has not been investigated on a transcriptional level. We provide evidence for a regulation of mitochondrial function by ethylene and auxin predominantly in the epidermis. The observed down-regulation of ethylene biosynthesis genes concomitant with reduced expression of genes encoding proteins of auxin biosynthesis and/or uptake in the epidermis contrasts with the mesophyll and vasculature. This may also link mitochondrial signalling to growth regulation, previously described on a whole plant level through the transcription factor ANAC017 (Meng et al., 2019), which might a have a component specific to the epidermis similar to shade avoidance (Procko et al., 2016). The auxin biosynthesis-promoting role of ethylene (Stepanova et al., 2005) also explains the overlapping transcriptional of responses to mitochondrial dysfunction, flooding and hypoxia (Wagner et al., 2018). An antagonistic relationship of mitochondrial retrograde signalling and auxin has been found before (Ivanova et al., 2014; Kerchev et al., 2014), and our results suggests an additional, tissue-specific layer for this regulation involving ethylene. This provides further evidence for the specialised roles of the leaf tissue types for stress responses.

In conclusion, our study determined the expression levels of Arabidopsis genes in the epidermis, mesophyll and vasculature, their specificity, and their changes in response to five stimuli that induce stress signalling pathways. Together these data provide a rich reference data set for further studies on the specific functions of leaf tissues and cells and their role for stress tolerance. Furthermore, networks of genes and their upstream regulators in each of the tissues were exemplary defined for the AA treatment, providing information on regulatory networks and promoter sequences.

The apparent absence of a common response to different stresses in the tissues might provide an advantage for the engineering of more stress tolerant crops as this allows a combinatorial stacking of the identified tissue-specific tolerance pathways tailored to the environmental challenges.

## METHODS

### Plant material and stress treatments

Seeds of *Arabidopsis thaliana* (accession Col-0) were surface sterilised, stratified for 2 d at 4°C in the dark and sown on Gamborg’s B5 Basal Salt medium (Sigma G5768) supplemented with 1% (w/v) sucrose, 0.75% (w/v) agar, B5 vitamins (Sigma G2519) and 0.05% (w/v) MES hydrate adjusted to pH 5.8. Seedlings were grown at 23°C in a 16 h light (100 μmol m^−2^ s^−1^) and 8 h dark photoperiod for 12 d. For stress treatments, plants were sprayed to run-off with solutions of 50 μM antimycin A, 10 mM 3-amino-1,2,4-triazole, 1 mM methyl viologen, 2 mM salicylic acid and water as the control treatment, and harvested after 3 h. For treatment with ultraviolet light, plants were exposed to an intensity of 1.5 W/m^2^ for 20 min before harvest.

### Laser capture microdissection

Briefly, after harvesting the first true leaves were immediately fixed in Farmer’s solution (3:1 ethanol:acetic acid) by infiltration for 10 min in a vacuum and after further incubation for 4 h processed using the Leica Semi-Enclosed Benchtop Tissue Processor TP1020 (Leica Biosystems) for an ethanol series (75 %, 85 %, 3 x 100 % (v/v) for 1 h each), followed by an ethanol: xylene series (75:25 %, 50:50 %, 25:75 % (v/v) for 1 h each) and a final xylene series (2 x 100 % (v/v) for 1 h each). Leaves were then embedded in Surgipath Paraplast^®^ Paraffin (Leica Biosystems). Sections of 10 μm thickness were prepared using a Leica Fully Automated Rotary Microtome RM2265 (Leica Biosystems), transferred to a Zeiss 1 mm PEN membrane-coated microscopy slide, and dried at 37°C for 30 min. Paraffin was removed by two washes with of 100 % xylene and sections air-dried. Laser dissection was performed with a PALM MicroBeam LCM system with Palmrobo v 4.6 software (Zeiss). Between 200 to 300 cuts from 5 sections were pooled for each biological replicate, corresponding to approximately 2000 cells for epidermis and mesophyll, respectively, and 6000 cells for vasculature. RNA from these cells was isolated immediately after cell isolation.

### RNA isolation and sequencing

For three biological replicates of each treatment and tissue, total RNA from captured cells was isolated with the PicoPure RNA isolation kit (Thermo Fisher Scientific, Australia) according to the manufacturer’s instructions. After removal of DNA using the on-column DNase digestion Set (Sigma-Aldrich Australia), RNA quality and quantity was assessed on a TapeStation 2200 system (Agilent). Next-generation sequencing libraries were prepared using an Ovation SoLo RNA-Seq System following the instructions of the manufacturer (Nugene) and sequenced on a NextSeq500 system (Illumina) as 70 bp single-end reads with an average quality score (Q30) of above 90% and an average of 6.7 M aligned reads per sample.

### Bioinformatic analyses

Read quality control was performed using the FastQC software (https://www.bioinformatics.babraham.ac.uk/projects/fastqc/). The abundance of transcripts and estimates of counts was quantified on a gene level by pseudo-aligning reads against a k-mer index build from the representative transcript models downloaded for the Araport 11 annotation (Cheng et al., 2017) using a k-mer length of 31 using the kallisto program with 100 bootstraps (Bray et al., 2016). The program sleuth with a Wald test was used to test for differential gene expression (Pimentel et al., 2017). Only genes with at least 5 counts in a quarter of all samples were included in the further analysis. Genes were called as differentially expressed with a |log_2_ (fold change)|>1 and a false discovery rate FDR < 0.05. For further analyses, hierarchical clustering and generation of heat maps the Partek Genomics software suite version 6.16 (Partek Incorporated, http://www.partek.com/) was used. GO term enrichment analysis was performed using the ClueGO plugin for Cytoscape (Bindea et al., 2009). Circular plots were generated with the circlize package in R (Gu et al., 2014). Differences in expression of genes in functional categories was performed using a Wilcox rank sum test and with a p < 0.05 after Bonferroni correction for statistical significance in the PageMan software (Usadel et al., 2006). RNA-seq read data were deposited at the NCBI SRA database under project ID PRJNA668247. For the gene regulatory network generation the TF2Networks tool was used (Kulkarni et al., 2018). Transcription factors predicted as up-stream regulators of tissue-specific or unspecific DEGs were retained with a cutoff of q < 0.01. The final network was constructed in Cytoscape 3.6.1 (Shannon et al., 2003) using a yFiles radial layout algorithm (yWorks, Germany).

## AUTHOR CONTRIBUTIONS

JW, YX and OB designed the research. YX, YW, LCL, YZ performed the experiments. OB analysed the data. OB, JW, MGL interpreted the data. OB and JW wrote the manuscript with contributions by all authors.

## ACKNOWLEDGEMENTS

This work was supported by the facilities of the Australian Research Council Centre of Excellence Program (CE140100008, ARC Centre of Excellence in Plant Energy Biology; JW).

We acknowledge the La Trobe University Genomics Platform for access to next generation sequencing equipment.

## Supplemental Information

Supplemental Figure 1.

## SUPPLEMENTAL INFORMATION

**Supplemental Figure 1:**
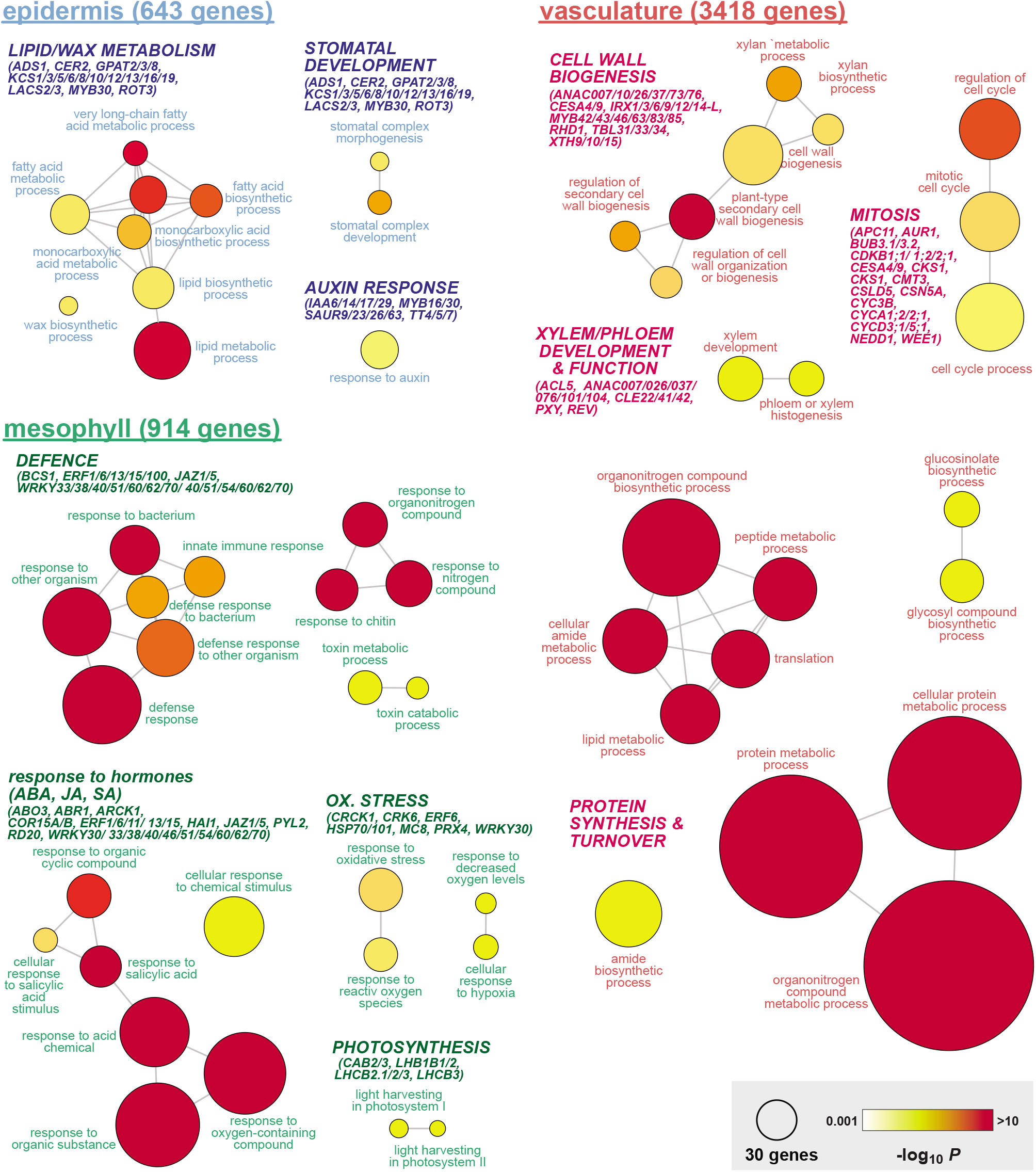
GO term enrichment analysis for genes with tissue-specific expression. GO term enrichment (p < 0.001 after Bonferroni correction) was analysed using the ClueGO plugin for Cytoscape (REF) for genes specifically expressed in one of three tissues as indicated (Supplemental Table 2). Circle size represents the number of genes asscociated with a GO term and circle colour the corresponding p-value. Key genes included in the corresponding GO terms are given in brackets.

**Supplemental Figure 2:**
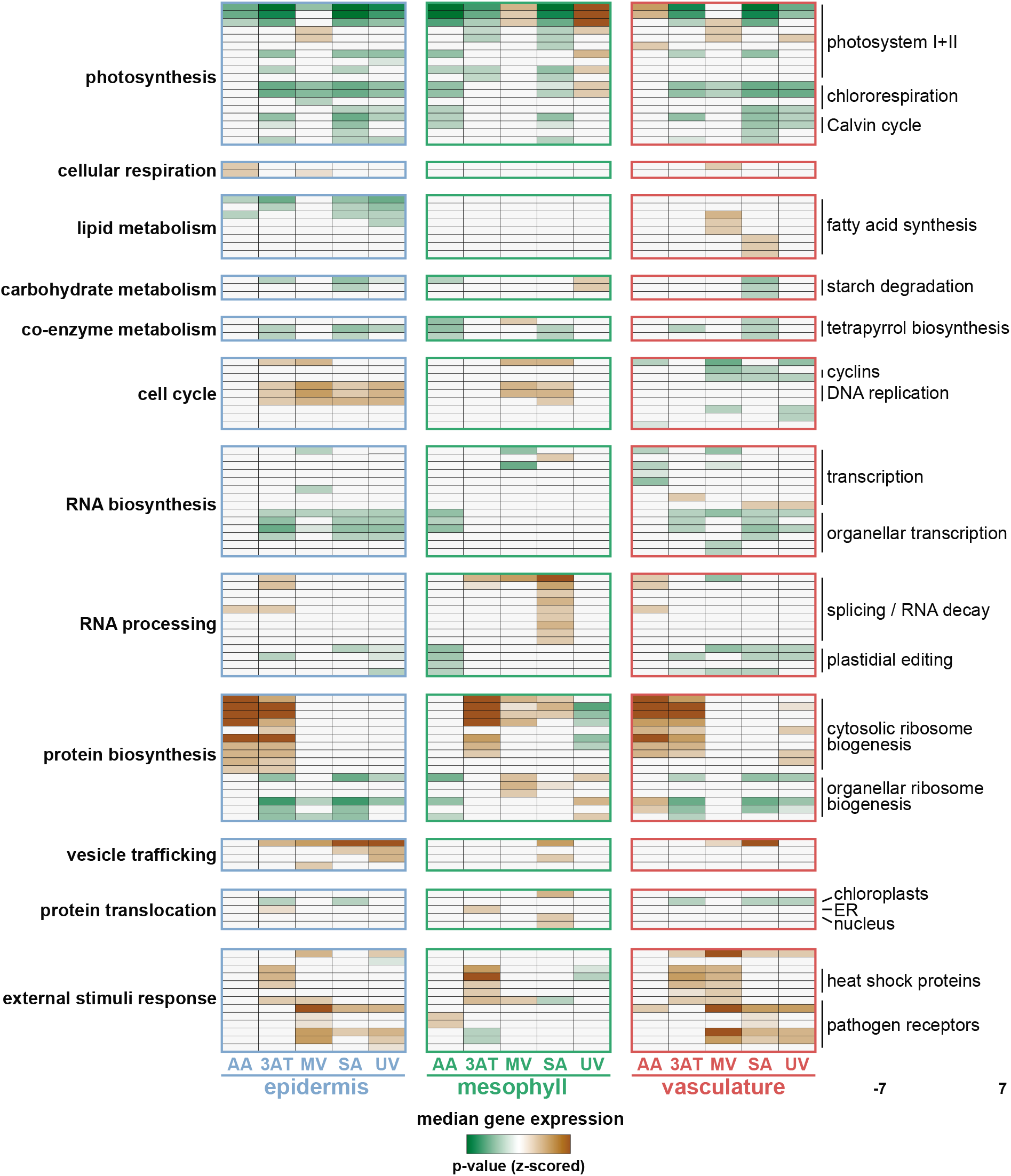
Overrepresentation analysis of DEGs in functional categories. Extended listing of overrepresented functional categories presented in Figure 2. Using the PageMan tool, the difference in expression of genes in functional categories was determined by comparison of the stress treatments with the control and is represented by heatmaps (Usadel et al., 2006). Indicated on the left are the annotations of high order bincodes and on the right selected, more detailed bincodes (see Supplemental Table 5 for complete listing). Colour scale indicates the p-value for statistical difference (Wilcox rank sum test, z-scored after Bonferroni correction) of the median fold-change of genes in the individual bins to all other genes. Brown and green colour indicate higher and lower median expression, respectively, and thus stronger stress responses of genes in these functional bins.

**Supplemental Table 1:** Expression of genes in three leaf tissues tissues under control conditions (given as transcripts per million, TPM) and their tissue specificity as determined by tau (Figure 1).

**Supplemental Table 2:** GO term enrichment analysis for genes specific for epidermis, mesophyll or vasculature (Figure 2). For each tissue, details for enriched GO term with the associated p-values after Bonferroni correction and the percentage of DEGs of the total number of genes within a given GO term.

**Supplemental Table 3:** List of tau values for genes in the different stress treatments and their tissue-specificity

**Supplemental Table 4:** Changes in tissue specificity. Given are percentages for the corresponding changes.

**Supplemental Table 5:** List of differentially expressed genes (DEGs) after stress treatments when compared to control conditions. DEGs were called with a |log_2_ (fold change)| > 1 and FDR < 0.05. Given are log_2_ (fold change) values.

**Supplemental Table 6:** PageMan analysis giving z-scored p-values of statistical difference of median expression of genes in the indicated functional bins to all other genes

**Supplemental Table 7:** List of DEGs in the intersects of the AA stress treatment (Figure 5A)

**Supplemental Table 8:** GO term enrichment analysis for genes specific for epidermis, mesophyll or vasculature (Figure 5B). For each tissue, details for enriched GO term with the associated p-values after Bonferroni correction and the percentage of DEGs of the total number of genes within a given GO term.

**Supplemental Table 9:** Regulator and target relationships for the gene regulatory network of DEGs in the antimycin A treatment as predicted by the T2Fnetwork tool.

